# Pectin remodeling belongs to a homeostatic system and triggers transcriptomic and hormonal modulations

**DOI:** 10.1101/2021.07.22.453319

**Authors:** François Jobert, Stéphanie Guénin, Aline Voxeur, Kieran J. D. Lee, Sophie Bouton, Fabien Sénéchal, Ludivine Hocq, Gaëlle Mongelard, Hervé Demailly, Petra Amakorová, Miroslav Strnad, Samantha Vernhettes, Gregory Mouille, Serge Pilard, J. Paul Knox, Ondřej Novák, Jérôme Pelloux, Laurent Gutierrez

**Author notes:** present address: Umeå Plant Science Centre, Swedish University of Agricultural Sciences, 90736 Umeå, Sweden (François Jobert). these authors contributed equally to this work (François Jobert, Stéphanie Guénin).

## Abstract

- Here, we focused on the biological modifications arisen from a strong and transient variation of the pectin methylesterification status during the seed-to-seedling transition.
- A reverse genetic approach was used to trigger specific reduction of pectin de-methylesterification during the seed maturation stage and the related physiological effects were assessed using a combination of biochemical, transcriptomic and microscopic analyses.
- Arabidopsis *PME36* is required to implement the characteristic pattern of de-methylesterified pectin in the mature seed. While this pattern is strongly impaired in *pme36-1* and *pme36-2* mature seed, no phenotypical effect is observed in the knockout mutant during seed germination. By analyzing hormone homeostasis and gene expression regulation, we show a strong and dynamic physiological disorder in the mutant, which reveals the existence of a complex compensatory mechanism overcoming the defect in pectin de-methylesterification.
- Our results reveal that pectin methylesterification status acts as upstream modulator involved in an undescribed homeostatic system in which pectin remodeling, hormone signaling and transcriptomic regulations interact to ensure the maintenance of a normal seed-to-seedling developmental program.

## INTRODUCTION

Plant cells are confined by semi-rigid walls consisting of a polysaccharide network, composed of cellulose, hemicellulose and pectins, which contains proteins and ions (Chebli *et al*., 2021). How cell wall plasticity is controlled to regulate plant development and growth is one of the outstanding questions in plant biology (Cosgrove, 2018). During the last decade, pectin has emerged as an important player in morphogenesis regulation, through its effects on wall stiffness modulation that is required for many developmental processes, among which seed development and germination (Müller *et al*., 2013; Levesque-Tremblay *et al*., 2015a; Scheler *et al*., 2015). Pectins, mainly composed of homogalacturonan (HG), are secreted into the wall in a methylesterified form – i.e. pectins with a high degree of methylesterification (DM) – and are then de-methylesterified *in muro* by the action of pectin methylesterase (PME) proteins, the activities of which are regulated by pectin methylesterase inhibitor (PMEI) proteins. For a long time, pectin de-methylesterification was considered to lead to only to cell wall stiffening, though the production of rigid “egg-box” structures resulting from calcium-mediated cross-linking of HG with low DM (Willats *et al*., 2001). Conversely, in recent years pectin de-methylesterification has been shown to result in cell wall softening, by a mechanism that remains elusive and in which pectin degradation enzymes – active on low DM pectins – were thought to play a role (Levesque-Tremblay *et al*., 2015b). Since then, the effect of pectin DM appears to be mostly dependent on the cellular context. In addition, recent insights into the complexity of pectin interactions with other cell wall components (Du *et al*., 2020), the existence of cell wall domains harboring spatially restricted pectin modifications (Francoz et al., 2019), and the proposition that pectin nanofilaments are involved in cell expansion (Haas et al., 2020), raise challenges to a full comprehension of the role of pectin methylesterification in cell wall plasticity during plant development.

Deciphering the control of pectin de-methylesterification appears even trickier. PME and PMEI proteins are encoded by multigenic families, composed of 67 and 76 genes in Arabidopsis, respectively (Wang *et al*., 2013; Hocq *et al*., 2017b). Efficiency of PMEI inhibitory activity depends on PME/PMEI pairings in a pH-depend manner (Hocq *et al*., 2017b; Sénéchal *et al*., 2017). Obviously, the high number of possible combinations restricts the possibility of *in planta* functional analysis of PME/PMEI interactions. Pectic HG de-methylesterification reactions drive acidification of the cell wall, whereas PME activity optima are at alkaline pH and inhibitory activity of PMEI is promoted at acid pH (Sénéchal *et al*., 2015, 2017; Hocq *et al*., 2017b). Thus, pectin de-methylesterification activity turns out to be dynamically regulated within cell walls, through feedback loops involving local pH changes (Hocq *et al*., 2017a), which adds a further level of complexity in the analysis of this enzymatic system.

Knockout mutations in *PME* genes have been reported to result in developmental defects in only a very limited number of studies and the absence of a phenotype observed in a single *pme* mutant is usually attributed to gene functional redundancy, although compensatory mechanisms between *PME* genes have never been extensively characterized. Most of the time, the focus is on the impact of overall pectin methylesterification modification, using plants overexpressing a *PME* or – more often – *PMEI* gene which triggers a steady dominant change in PME activity. Although too drastic to allow the analysis of the fine-tuned control of pectin de-methylesterification, such approaches seem appropriate to study the role of pectin methylesterification from a mechanical perspective. Nevertheless, beyond their structural role, pectins and pectin-derived degradation products, such as oligogalacturonides (OGs), interact with wall sensors and are particularly prone to trigger signaling pathways involved in the cell wall integrity (CWI) maintenance (Wolf, 2017; Vaahtera *et al*., 2019). On the one hand, pleiotropic effects of *PME*/*PMEI* gene overexpression brings about loss of pectin remodeling dynamics and overrides the subtle effects it could produce, especially regarding this signaling function. On the other hand, absent or subtle phenotypes of *pme* mutants have never really encouraged in-depth investigations of possible modulations of their physiological parameters. Consequently, our knowledge of the physiological effects resulting from transient modification of pectin methylesterification remains poor.

In cell wall mutants, the discrimination between the direct effects of the mutation on wall composition and those indirectly arising from signaling-related responses is always hard. However, a few clues indicate that some important signaling pathways could be activated by pectin methylesterification status during plant development. Auxin is known to regulate the DM of pectins (Voiniciuc *et al*., 2013; Schoenaers *et al*., 2018) and, reciprocally, when HG de-methylesterification is inhibited, auxin signaling is altered in the shoot apical meristem by modification of the localization of the auxin transporter PIN1 (Braybrook & Peaucelle, 2013). More recently, pectin methylesterification was also shown to impact on auxin transport during apical hook formation, suggesting the existence of a feedback loop between auxin and HG (Jonsson *et al*., 2021), the molecular basis of which remains unknown.

In this context, we set out to determine the extent to which physiological modifications could originate from impairment in pectin methylesterification status, and to determine the impact this potential signaling function of pectin could have on plant development. For this purpose, we used the knockout *pme36-1* mutant line, deficient for *PME36* gene expression, in which pectin methylesterification status was strongly impaired in mature seeds. Interestingly, we discovered that several regulatory pathways were affected in the mutant, including hormone signaling and transcriptomic regulation, demonstrating that disruption in pectin methylesterification pattern can directly trigger large physiological changes. Surprisingly, concomitantly to this dynamic physiological disorder, the mutant followed a germination program which was strictly the same as in the wild type. This led us to hypothesize that the physiological changes observed in *pme36-1*, far from being a disruption, actually revealed the existence of a homeostatic system connecting different regulatory pathways, which ensure the maintenance of the seed-to-seedling transition. This system was acting in the *pme36-1* mutant by adjusting different physiological parameters, thereby restoring a new equilibrium suitable for maintaining normal developmental process in the mutant background.

## MATERIALS AND METHODS

### Plant material and growth conditions

The Arabidopsis *pme36-1* and *pme36-2* mutants were isolated from the SALK T-DNA collection (SALK_022170 and SALK_091350 respectively, Col-0 ecotype). The left flanking sequence of the T-DNA insertion sites were amplified by PCR on genomic DNA using the T-DNA specific primer LB (5’-CGATTTCGGAACCACCATCAAACAGGA-3’) combined to *AtPME36* specific primers KOF (5’-ACGGGACATAACGTTCGAGA-3’) or KOR (5’-TCGATTTTCCTCCCTTTCG-3’). PCR products were sequenced and the insertion in SALK_091350, although predicted to be in the third exon, was actually identified as being in the third intron (Fig. S1). Both mutant lines were characterized as knockout for *PME36* gene expression by RT-PCR using primer pair flanking the insertion sites (Fig. S1). *TIP41* amplicon (Gutierrez *et al*., 2012) was used as RT-PCR control in Fig. S1C and as reference to normalize RT-qPCR data in Fig. S1D. Plants were grown on soil in a greenhouse under the following conditions: 16 h light (120 µmol m^−2^ s^−1^, 20°C)/8 h dark (15°C) cycles.

### Germination testing

Mature dry seeds were stored at room temperature for three months after harvesting and were then sterilized and sown in vitro as previously described (Gutierrez *et al*., 2009). Germination was analyzed under long-day conditions (16h/8h) at 20°C. *In vitro* plates were transferred to the light to induce germination, which constitutes the time point 0 h. Time measurement was defined as hour after induction (HAI), for seeds transferred just after sowing, and as hour post stratification (HPS), for seed transferred after a previous 48 h-period in dark at 4°C for stratification. Germination was scored with a SteREO Discovery V20 stereomicroscope (Zeiss). Testa and endosperm ruptures were scored as in Müller et al. (2013). The effects of different hormones on germination were also assessed by adding 5 µM of abscisic acid (ABA) or 10 µM of auxin (IAA) (Sigma) in the medium. For each biological replicate, at least 50 seeds were analyzed and values in the graph are the mean from 3 biological replicates.

### Analysis of promoter activity

Cloning of one kb of the *PME36* promoter, plant transformation and selection, and β-glucuronidase staining, were carried out as previously described (Guénin *et al*., 2011). Plant samples were destained in 75% (v/v) ethanol and images were acquired using a SteREO Discovery V20 stereomicroscope (Zeiss).

### RNA Isolation and cDNA Synthesis

RNAs from *pme36-1, pme36-2* and Col-0 siliques, seeds and seedlings were prepared as previously described (Gutierrez *et al*., 2006).

### Real-time RT-PCR analysis

Transcript levels were assessed by quantitative RT-PCR using a LightCycler 480 system (Roche), as described by Gutierrez *et al*. (2012). Primers sequences for *PME* and *PMEI* transcripts (Table S1), designed towards the 3’ end of the RNA sequence using QuantPrime (https://quantprime.mpimp-golm.mpg.de/) (Arvidsson *et al*., 2008) and Primer3 software, were validated by melting curve analysis and efficiency measurement.

### Affymetrix microarrays analysis

16 and 24 h post-stratification seeds were collected, flash frozen and ground into liquid nitrogen. Total RNA extraction, RNA quantification and quality control, labelled single-stranded DNAs production, hybridization on Affymetrix four-arrays strips (Arabidopsis Gene 1.1 ST Array strip), imaging and data processing were performed as in Voxeur *et al*. (2019). Normalized expression values were filtered for statistical relevance of differential expression using FDR F-Test p-value<0,01 and are shown in Dataset S1. Affymetrix Microarray data are available in the Gene expressionOmnibus database (https://www.ncbi.nlm.nih.gov/geo/query/acc.cgi?acc=GSE155433)

### Enzymatic fingerprinting of pectins

Alcohol-insoluble material from ground seeds or hypocotyls was incubated during 16h at 37°C in 50 mM ammonium formate pH 5 with 5 units of *Aspergillus aculeatus* endo-polygalacturonase (Megazyme), which hydrolyzed the HG between at least two non-esterified GalA residues and was tolerant to acetyl groups (Voxeur *et al*., 2019). The digestion products (oligogalacturonides, OGs) were separated according to their degrees of polymerisation and methylesterification using size exclusion chromatography (SEC) followed by mass spectrometry analysis, as previously described (Hocq *et al*., 2020).

### Extraction of protein fraction and determination of PME activity

Samples were ground in liquid N_2_ to obtain a fine powder and then transferred in the extraction solution. Proteins were extracted, quantified and assayed for total PME activity as in Guenin *et al*. (2001). Data are the means of 9 independent replicates and were statistically analyzed by a Mann-Whitney test (Statisca).

### Immunofluorescence microscopy

Dry seeds were fixed, dehydrated and infiltrated with LR White resin (London Resin Company) as previously described (Lee *et al*., 2012). Sections were cut to a thickness of 0.5 μm using a diamond knife on an Ultracut microtome (Reichart-Jung) and collected on multiwell slides (ICN Biomedicals) coated with Vectabond reagent (Vector Laboratories). Calcofluor staining and immunolocalisation with anti-HG antibodies JIM7 and LM20 was conducted as previously described (Lee *et al*., 2012). Epifluorescence and light microscopy analyses were performed on cross sections from at least three seeds.

### Quantification of phytohormones

50 mg of fresh weight mature seeds, 24HPS and 48 HPS seedlings were harvested, dried on tissue paper and frozen in liquid nitrogen. The samples were prepared in three biological replicates and phytohormones were extracted using an aqueous solution of methanol (10% MeOH/H_2_O, v/v) from 1 mg of samples. The concentrations of endogenous auxin metabolites (free IAA, its conjugates IAAsp and IAGlu, and catabolite oxIAA), and ABA were determined in three independent experiments, each composed of three biological replicates. A cocktail of stable isotope-labelled standards was added, 5 pmol of [^13^C_6_]IAA, [^13^C_6_]oxIAA, [^2^H_5_][^15^N]IAAsp, and [^2^H5][^15^N]IAGlu and 10 pmol of [^2^H_6_]ABA (all from Olchemim Ltd) per sample to validate the LC-MS/MS method using stable isotope dilution method. The extracts were purified using Oasis HLB columns (30 mg/1 ml, Waters) and targeted analytes were eluted using 80% MeOH according to the method previously described (Floková *et al*., 2014). Separation was performed on an Acquity UPLC™ System (Waters) equipped with an Acquity UPLC BEH C18 column (100 × 2.1 mm, 1.7 μm; Waters), and the effluent was introduced into the electrospray ion source of a triple quadrupole mass spectrometer Xevo™ TQ MS (Waters).

## RESULTS & DISCUSSION

### PME36 is required to establish the demethylesterification pattern of HG in mature seeds

Using RT-qPCR analysis in Col-0 plants and GUS staining of the promoter-reporter line, we assessed the expression pattern of the *PECTIN METHYLESTERASE 36* (*PME36*) gene during seed maturation and early seedling development. The transcript level peaked in siliques from 13 to 16 days post-anthesis (DPA) (**Fig. 1A**). A pool of *PME36* mRNAs was stored in the mature seed, which rapidly decreased in 24 and 48 hours post-stratification (HPS) seedlings. In seeds and in 24 HPS seedlings, the promoter displayed constitutive activity, whereas in 48 HPS seedlings the GUS staining appeared stronger in cotyledons and in the upper part of the hypocotyl, indicating that residual gene expression was restricted to these organs at this latter stage (**Fig. 1B**).

**Figure 1.**
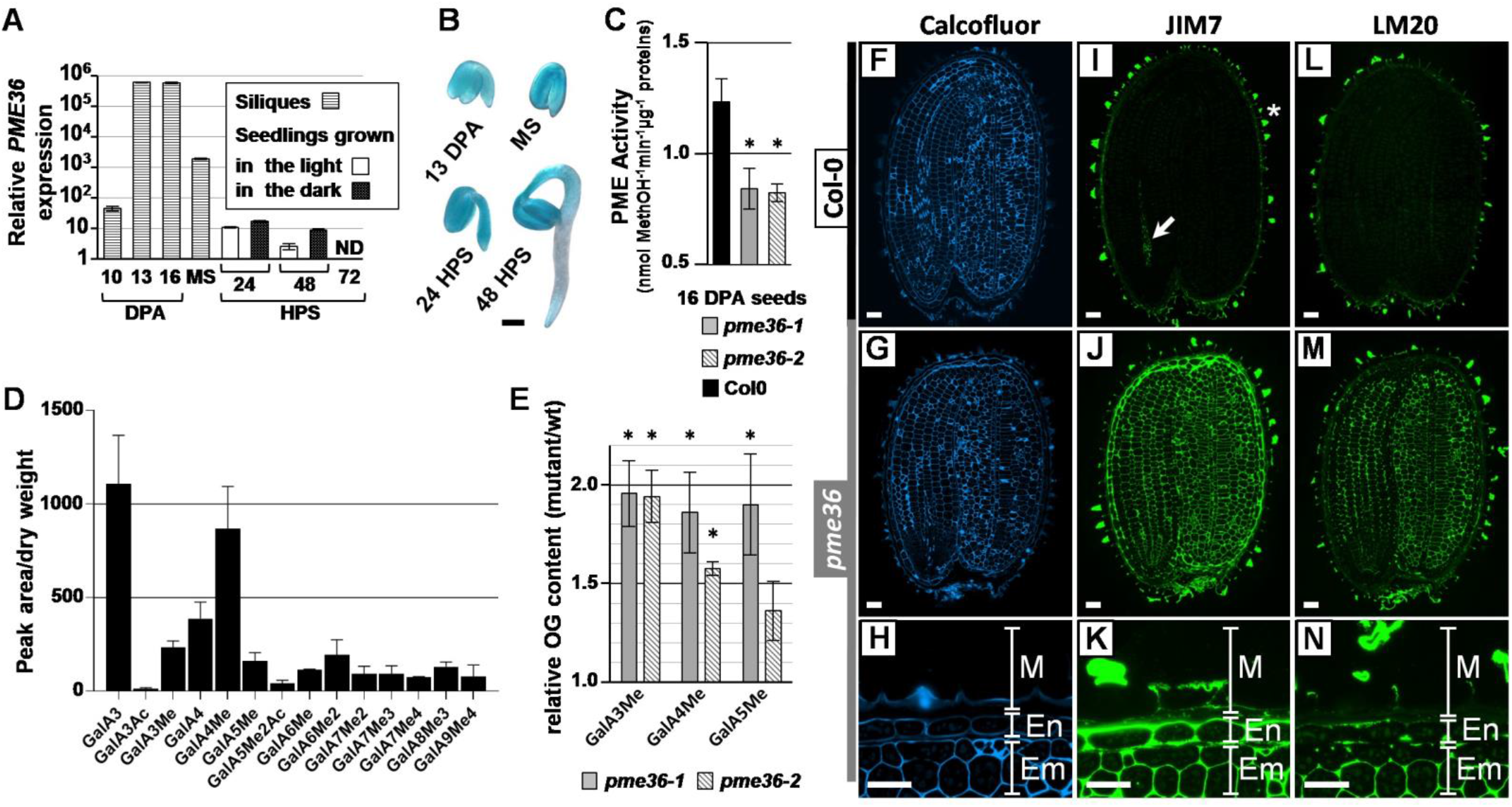
*PME36* is required to establish the demethylesterification patern of HG in mature seeds. **(A)** Quantification by RT-qPCR of *PME36* transcripts in 10, 13 and 16 days post anthesis (DPA) Col-0 siliques, in mature seeds (MS) and in 24 and 48 hours post stratification (HPS) Col-0 seedlings grown in long-day conditions or in the dark. In 72 HPS Col-0 seedlings, no *PME36* transcripts were detected whatever the growth conditions. Error bars indicate +/- SE obtained from 3 qPCR replicates. **(B)** GUS staining of *promPME36:GUS* in 13 DPA seeds, in mature seeds (MS) and in 24 and 48 HPS seedlings grown in the dark. Bar = 200 µm. **(C)** PME activity was assessed in Col-0, *pme36-1* and *pme36-2* 16 DPA seeds. A Mann-Whitney Comparison Test indicated that PME activity was significantly reduced (*) in 16 DAF seeds from both mutants compared to the wild-type (P<0.05). Error bars indicate SE obtained from three independent biological replicates, each one composed of three technical replicates. **(D)** Enzymatic fingerprinting of pectins from mature Col-0 seed. **(E)** Relative increase of OG content, produced by enzymatic fingerprinting of pectins, in the mature *pme36-1* and *pme36-2* seed compared to the wild type for which the level is set to 1. Sidak’s multiple comparisons test determined that significantly higher quantities (*) of GalA3Met, GalA4Met and GalA5Met were released from *pme36-1* and *pme36-2* than from Col-0 seed. **(F)** to **(N)** Medial sections of resin-embedded Col-0 and *pme36-1* mature seeds stained by Calcofluor White (F-H), which reveals all cell walls, and labelled with monoclonal antibodies JIM7 (I-K) and LM20 (L-N), which target HG with specific patterns and degrees of methylesterification. (H), (K), (N) show magnification of a part of the seed tegument from (G), (J), (M) sections, respectively. **(I)** In Col-0 seeds, partially methylesterified HG recognised by JIM7 was restricted to mucilage (*) and root vascular tissue (arrow). **(J)** and **(K)** The JIM7 epitope was abundant in embryo, endosperm and mucilage of *pme36-1* seeds. **(L)** The LM20 epitope was restricted to the seed testa/mucilage in Col-0. **(M)** and (**N)** The LM20 epitope was present in the embryo, mucilage and, at lower levels, endosperm of *pme36-1* seeds. Em: embryo; En: endosperm; M: mucilage. Scale bars = 25 µm.

The high expression of *PME36* during maturation suggested that the encoded protein might play an important role in the control of HG methylesterification in the seed. To address this question, we identified the *pme36-1* and *pme36-2* mutants, two T-DNA insertion lines that were knockout for *PME36* gene expression (**Fig. S1**). PME activity was significantly reduced in 16 DPA seeds from both mutants compared to the wild type, indicating the importance of PME36 to the total PME activity during seed maturation (**Fig. 1C**). In order to analyze whether the reduction of PME activity during seed maturation led to a concurrent increase in the level of methylesterified HG in mature seed from both mutants, we analyzed the methylesterification of pectins by enzymatic fingerprinting in mature seeds from *pme36-1, pme36-2* and Col-0 (Voxeur *et al*., 2019; Hocq *et al*., 2020). This technic allowed to provide the seed-specific OG profile obtained from HG hydrolysis by endo-polygalacturonase in Col0 mature seed (**Fig. 1D**). In the mutants, differences in OG profiles revealed that more GalA3Me, GalA4Me and GalA5Me were released from *pme36-1* and *pme36-2* than from Col-0 seeds, suggesting the presence of higher number of hydrolysable blocks of methylesterified HG (**Fig. 1E**). To characterize the pattern of HG methylesterification within the seed, sections of resin-embedded mature seeds from *pme36*-1 and Col-0 were probed with two different monoclonal antibodies directed to HG with specific patterns and degrees of methylesterification (DM). The antibody JIM7 was previously shown to display affinity for HG with medium DM and LM20 was characterized as preferentially binding to HG with medium to high DM (Clausen *et al*., 2003; Verhertbruggen *et al*., 2009). Labelling with Calcofluor White highlighted cell walls in medial longitudinal sections of Col-0 and *pme36-1* mature seeds (**Fig. 1F,G**). Higher magnification views of the *pme36-1* sections allowed analysis of seed mucilage [M], endosperm [En] and embryo tissues [Em] (**Fig. 1H**). JIM7 labelling revealed that HG with medium DM was restricted to the mucilage and the root vascular tissue in the wild-type mature seeds (**Fig. 1I**). On the contrary, the JIM7 epitope was widely distributed in *pme36-1* seeds, where it was abundant in embryo and endosperm as well as in the mucilage cells (**Fig. 1J,K**). HG epitopes recognized by JIM7 are known to be partially sensitive to polygalacturonase treatment (Willats *et al*., 2000). The higher amount of GalA3Me, GalA4Me and GalA5Me released by the PG is therefore consistent with the increased JIM7 labelling observed in *pme36-1* mature seed (**Fig. 1E**). The LM20 epitope was only detected in the seed mucilage in Col-0, whereas it was present in the embryo, the mucilage and, to a lower extent, the endosperm of *pme36-1* seeds, confirming the presence of HG of medium to high DM in the cell walls of mutant mature seeds (**Fig, 1L,M,N**). Altogether, these results showed that, while Col-0 mature seeds contained a relative small amount of methylesterified HG, *PME36* defect strongly impaired this pattern with substantial enrichment in HG with medium to high DM in the mutant mature seeds. This indicates the crucial role PME36 plays in the orchestration of the demethylesterification of pectins in mature seeds of Arabidopsis.

### Increased pectin methylesterification in *pme36-1* seed is rapidly rectified and does not affect the germination process

In Arabidopsis *PMEI5* overexpressing seed, in which PME activity was strongly reduced, cell wall loosening was induced by the increase of pectin DM, which led to faster seed germination by facilitating testa rupture and radicle emergence through the endosperm (Müller *et al*., 2013). As a consequence, PME activity seems not to be necessary *per se* to achieve the germination process. However, specific regulation of PME activity was observed in Arabidopsis germinating seeds, peaking during testa rupture and then declining once endosperm rupture was reached. Moreover, analysis of testa rupture in garden cress seed added complexity into this model, highlighting specific regulations of PME expression occurring in different seed compartments, and also the antagonistic effects of PME treatment on testa rupture, which can be either positive or negative depending on the concentration of enzyme applied (Scheler *et al*., 2015). Altogether, these studies showed the crucial – although ambiguous – role of PME-related regulation of pectin DM in the germination process. In consequence, *pme36-1* seeds were expected to germinate faster than the wild type, given their strong enrichment in methylesterified pectins, as in *PMEI5* overexpressing seeds. We assessed the germination kinetics by determining the timing of testa and endosperm ruptures in post-stratification Col-0 and *pme36-1* seeds. Surprisingly, the germination kinetic was the same in the mutant as in the wild type (**Fig. 2A**). This unexpected result suggested that the strong defect in pectin methylesterification pattern observed in *pme36-1* mature seed might have disappeared very quickly during the early germination stage, i.e. before testa rupture. To investigate this hypothesis, first we assessed the PME activity in germinating seeds from mutant and wild type. In 16 HPS seeds, before testa rupture stage, the PME activity in *pme36-1* was already reestablished to the wild-type level, and in 40 HPS seed, when endosperm rupture stage was reached, the increase in PME activity level was the same in the mutant as in the wild type (**Fig. 2B**). This result indicated that the regulation of PME activity was fully restored in *pme36-1* seed at very early stage of germination. The remaining question was to determine whether this recovery led to subsequent restoration of normal DM of pectin in the young seedling. To answer this, we analysed the methylesterification of pectin by enzymatic fingerprinting in 48 HPS mutant and wild-type seedlings. Despite the increase in GalA3 and GalA4Me previously observed in *pme36-1* mature seed, in 48 HPS seedlings a similar OG profile was obtained for the two lines, indicating the mutant recovered a wild-type level of pectin DM during germination (**Fig. 2C**). Overall, our results showed that, although *pme36-1* mature seed displayed strong alterations in pectin methylesterification, the mutant rapidly regained a normal and steady ability to regulate pectin demethylesterification, before the testa rupture stage of germination. This led *pme36-1* to recover a normal pectin methylesterification pattern and thus normal testa and endosperm ruptures, thus avoiding pectin-related effects on germination.

**Figure 2.**
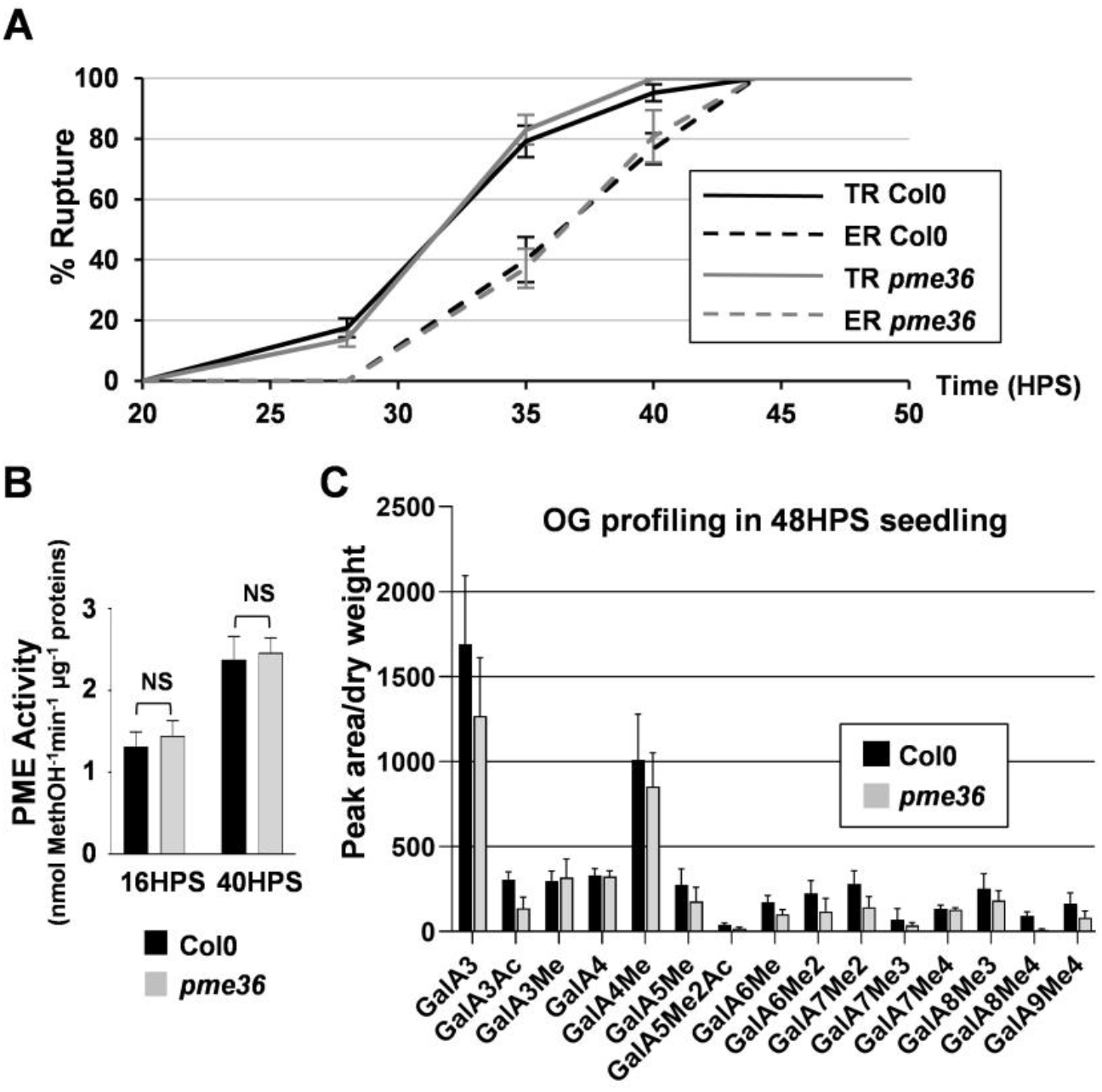
Impaired pectin demethylesterification in *pme36-1* seed is rapidly rectified and does not affect the germination process. **(A)** Seed germination kinetic of *pme36-1* and Col-0 seeds characterized by testa rupture (TR) and endosperm rupture (ER). Data represent averages and error bars indicate +/- SE, obtained from three independent biological replicates of at least 50 seeds each. **(B)** PME activity was assessed in Col-0 and *pme36-1* 16 and 40 HPS seeds. A Mann-Whitney Comparison Test indicated that PME activity was not significantly different in *pme36-1* compared to the wild-type (P<0.05). Error bars indicate SE obtained from three independent biological replicates, each one composed of three technical replicates. **(C)** Enzymatic fingerprinting of pectins from 48 HPS Col-0 and *pme36-1* seedling, showing no difference between the two lines.

### Levels of *PME* and *PMEI* transcripts are strongly downregulated in *pme36-1* mature seed

The fast recovery in *pme36-1* of PME activity to the wild-type level at a very early germination stage, i.e. in 16 HPS seed, led us to hypothesize the involvement of transcriptional regulation of *PME* and/or *PMEI* genes in the mutant during the seed-to-seedling transition. Given the high sequence redundancy among both gene families and the possible variety of gene expression levels to be assessed, we designed a dedicated RT-qPCR assay allowing specific and sensitive quantification of transcripts from each of the 67 *PME* and 76 *PMEI* genes. By this means, quantification of individual *PME* and *PMEI* transcripts was performed in 13 DPA siliques, in mature seeds and in 24 HPS seeds from Col-0 and *pme36-1* lines (**Fig. 3**). The expression values shown for Col-0, represented with a red scale, are normalized non-calibrated RT-qPCR data, providing transcript levels that can be directly compared among genes. The relative expression values shown for *pme36-1*, are normalized RT-qPCR data that were calibrated to the wild-type level which was set to 1 (*i*.*e. pme36-1*/Col-0 expression ratio), providing fold-change levels of gene expression in the mutant compared to the wild type. This highlights the transcriptomic regulations triggered in the mutant, for which intensities are shown using a logarithmic bicolor scale, in red for upregulation and in blue for downregulation. Empty cells indicate absence of quantifiable transcripts. Transcript profiling of *PMEI* genes was very informative (**Fig. 3A**). In Col-0 13 DPA siliques, 50 different *PMEI* genes were expressed, among which 46 remained expressed in the mature seed, where 20 maturation-specific transcripts were also produced. Therefore, mature seed turned out to contain a very rich pool of *PMEI* transcripts, from 66 out of the 76 *PMEI* genes identified in Arabidopsis. By contrast, the diversity of *PMEI* transcripts strongly decreased in 24 HPS seed, where only transcripts from 30 genes remained produced, among which there were no new specific genes. Translation from stored mRNAs has been shown for a long time as essential in the seed germination process and constitutes a major mechanism of protein production during early germination (Rajjou *et al*., 2004; Bai *et al*., 2020; Sano *et al*., 2020). Therefore, the importance of the pool of *PMEI* transcripts found in mature seeds, followed by the fast decay of this mRNA pool in 24 HPS seeds, strongly suggested the necessity for a high diversity of PMEI proteins to be transiently produced during seed imbibition. These PMEI presumably induce inhibition of PME activity during imbibition, which could be required to activate the germination process at the cell wall level. This hypothesis is supported by the positive impact *PMEI5* overexpression played on germination activation (Müller *et al*., 2013).

**Figure 3.**
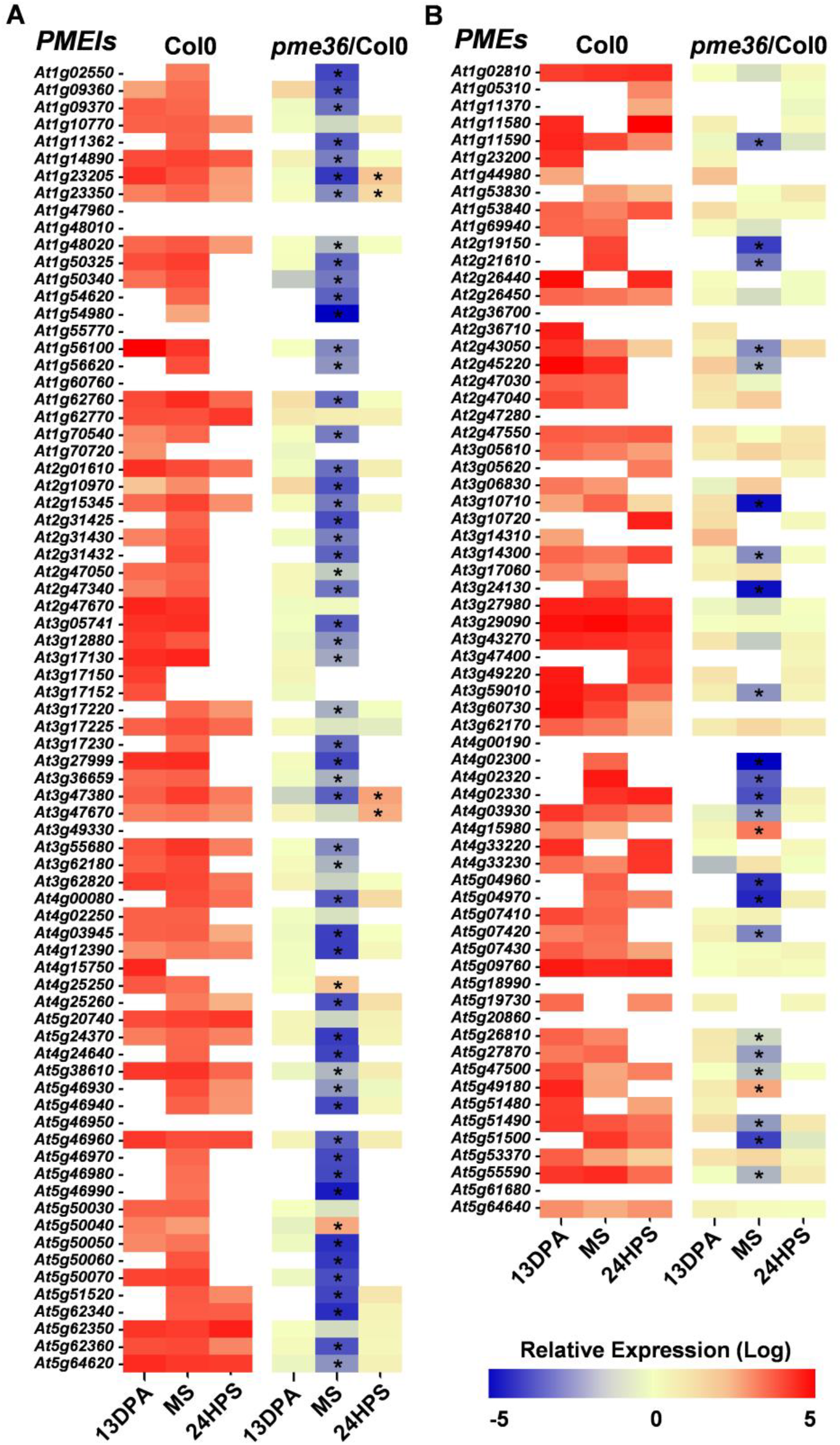
Levels of *PME* and *PMEI* transcripts are strongly downregulated in *pme36-1* mature seed. Quantification by RT-qPCR of *PMEI* (**A**) and *PME* (**B**) transcripts in 13 days post anthesis (DPA) siliques, in mature seed (MS) and in 24 hours post stratification (HPS) seedlings from Col-0 and *pme36*-1. Data in Col-0 are normalized non-calibrated gene expression levels (i.e. relative to the reference gene) and data in *pme36-1* are normalized data calibrated to the expression levels in the wild type which are set to 1 (i.e. *pme36-1*/Col-0). Transcript levels are shown using a logarithmic bicolor scale, in red for upregulation and in blue for downregulation. Empty cells indicate absence of quantifiable transcripts. Values are means from three independent biological replicates. A one-way analysis of variance combined with the Dunnett’s comparison posttest confirmed that the differences between the wild type and the mutants (*) are significant (P < 0.001, *n* = 3).

In 13 DPA siliques, the expression levels of *PMEI* genes were not significantly different in *pme36-1* compared to Col-0, indicating the absence of specific transcriptional regulation in the mutant at this stage (**Fig. 3A**). On the contrary, the level of 54 out of the 66 expressed *PMEI* transcripts dramatically decreased in the *pme36-1* mature seed, leading to a strong reduction in the *PMEI* transcript pool compared to the wild type. This *PMEI* transcriptional regulation was transitory, since expression of every gene returned to the wild-type level in 24 HPS *pme36-1* seed, except for 4 of them, for which expression slightly increased. This slight overexpression, as for 2 other *PMEI* genes in the mutant mature seed, was likely to be due to homeostatic adjustment or a specific role for these *PMEIs*, in response to the major alteration in *PMEI* gene expression occurring in *pme36-1* mature seed. The fact remained that the main effect of *PME36* disruption was a massive drop-down of *PMEI* transcript pool in the mutant mature seed, just as this pool raised its maximum in quantity and diversity in the wild type. Such a massive downregulation of gene expression is likely to trigger reduction of PMEI protein production at the onset of germination, thereby leading to increased PME activity by immediate release of PMEI-mediated PME inhibition. This would constitute an efficient compensatory mechanism that would explain by itself the fast reestablishment of PME activity and pectin methylesterification pattern to wild-type levels in *pme36-1* germinating seed (**Fig. 2B,C**).

Transcript profiling analysis provided a very different picture for *PME* genes (**Fig. 3B**). While the expression level in Col-0 was constant for a few genes from 13 DPA siliques to 24 HPS seeds, most of *PME* genes displayed transcriptional regulation that was very specific to each developmental stage. This revealed a much more dynamic transcriptional regulation for *PME* than for *PMEI* genes during the seed-to-seedling transition. In 13 DPA siliques and 24 HPS seeds, no difference in *PME* transcript level was observed in *pme36-1* compared to the wild type. On the contrary, the transcript level of 22 out of 45 expressed genes dropped in *pme36-1* mature seeds. Thus, like for *PMEIs*, but to a lower extent, the pool of *PME* transcripts decreased in *pme36-1* mature seeds compared to the wild type. While the decrease of *PMEI* transcript levels in *pme36-1* mature seeds appeared to be consistent with the compensation of the PME activity defect, the concomitant decrease of *PME* transcript levels in the mutant was counterintuitive. Interestingly, a similar counteractive mechanism was recently described in the control of Arabidopsis gynecium development by ETTIN transcription factor (Andres-Robin *et al*., 2018). ETTIN induced PME activity by negatively regulating the expression of *PMEI* genes, thus decreasing the DM of pectin, which caused cell wall loosening required to promote gynecium development. However, at the same time, ETTIN also negatively regulated the expression of several *PME* genes (Andres-Robin *et al*., 2020). The authors hypothesized this simultaneous negative regulation of *PME* expression, the significance of which appeared unclear in their model, to be a part of a “gas and brake mechanism”, in which antagonistic *PMEI* and *PME* genes were regulated in the same way in order to prevent excessive downstream effects. Our results confirm that such a mechanism is likely to exist and, by providing exhaustive expression data from both gene families, allow us to gain insight into the extent to which the transcriptional regulations of *PME* and *PMEI* genes might be connected.

Altogether, our results revealed the existence of a transcriptomic compensation in *pme36-1* seeds, consisting on specific regulation of *PME* and *PMEI* gene expression during the maturation stage. This would trigger, at the time of germination, a fast recovery of PME activity and pectin methylesterification pattern to the wild-type level in the mutant. Since this compensation necessarily needs an upstream activation and should be fine-tuned to adapt pectin DM to the germination requirements, we postulated the existence of a more complex homeostatic system which involved other regulations ensuring a normal germination process in *pme36-1* seeds. Such a system most likely requires crosstalk between different regulatory pathways, among which transcriptional regulation and hormone homeostasis are presumable players and for which further investigations were conducted in the *pme36-1* mutant.

### *pme36-1* displays dynamic gene expression regulation during germination

Two transcriptional phases were previously characterized during Arabidopsis seed germination, separated by the testa rupture stage and displaying distinct transcriptome changes (Dekkers *et al*., 2013). In order to identify potential transcriptional regulations specifically occurring during *pme36-1* germination, a transcriptomic analysis was performed in mutant and wild-type 16 and 24 HPS seeds, which characterized the stages of non-rupture seed imbibition and testa rupture, respectively (**Fig. 2A**). We identified a set of 374 genes differentially expressed in the mutant compared to the wild type (**Dataset S1**). Interestingly, while the developmental stages were spaced by only 8 hours, very few of the genes displayed the same transcription modulation (up- or down-regulation) in both conditions. The overwhelming majority showed either differential expression at only one stage or opposite regulation trends from one stage to another. This dynamic regulation exemplifies the specificity of the transcriptome at each transcriptional phase.

Three genes encoding polygacturonase (PG) proteins, among which *PGAZAT* that is involved in reproductive organ abscission in connection with abscisic acid (ABA) biosynthesis promotion (Ogawa *et al*., 2009; Xu *et al*., 2020), were deregulated in *pme36-1* germinating seeds. PGs lead to cell wall loosening by catalyzing the degradation of pectic HG with low DM. In order to restore a normal level in germinating *pme36-1* seeds, the DM of pectin necessarily decreased in the mutant from a very high level in the dry seed, to a wild-type level in 24 HPS seed. Interestingly, *PG* expression was upregulated in 16 HPS mutant seed, where pectins were probably still highly methylesterified, thus making them resistant to the lytic action of PGs. Then, in 24 HPS mutant seed, when pectins recovered a wild-type DM level and thereby an increased sensitivity to the degradation by PGs, *PG* expression was downregulated. This transcriptional modulation suggested the possible existence of a regulatory process fine-tuning PG level according to the DM of pectin. Additional cell wall-related genes were differentially expressed in the mutant during germination, such as *XTH4, XTH9* and *XTH25*, which are involved in xyloglucan metabolism. Xyloglucans are cell wall components and their impairment can impact seed germination (Sechet *et al*., 2016; Shigeyama *et al*., 2016).

Several genes related to ABA signaling were deregulated in *pme36-1*. First, *PYL13*, which is a member of the *PYL* family of ABA receptors (Fuchs *et al*., 2014), and *OST1*, encoding an ABA-activated kinase belonging to the SNF1-related protein kinases (SnRK2) family (Vlad *et al*., 2009), were strongly downregulated in 16 HPS *pme36-1* seeds. Both proteins being effective up-regulators of ABA signaling pathway, their down-regulation most likely led to decreased ABA signaling during *pme36-1* seed imbibition. Moreover, the *NCED9* gene, involved in ABA biosynthesis during seed development (Lefebvre *et al*., 2006), as well as *PER1* that reduces seed germination by suppressing the ABA catabolism (Chen *et al*., 2020), were both downregulated in 16 HPS *pme36-1* seeds, which might have an additive effect in decreasing ABA signaling through diminution of its levels in the mutant. A series of genes encoding typical ABA-induced seed proteins - like LEA, oleosin, oleosin-like, albumin and cruciferin proteins - were concurrently downregulated in 16 HPS mutant seeds, corroborating the possibility of ABA signaling reduction during mutant seed imbibition. However, at the same time, *HAI3* that encodes a PP2C ABA coreceptor inhibiting ABA signaling (Tischer *et al*., 2017) was down-regulated and *RAP2-6*, which overexpression has been shown to confer hypersensitivity to ABA signaling in seed germination (Zhu *et al*., 2010), was strongly up-regulated. Furthermore, the *SASP* gene, encoding a subtilase protein involved in reproductive development (Martinez *et al*., 2015) and in the regulation of ABA signaling (Wang *et al*., 2018), was also up-regulated in 16 HPS mutant seeds. Finally, *FAR1*, which positively regulates the ABA pathway (Wang & Wang, 2015), was first up-regulated in 16 HPS *pme36-1* seed and then down-regulated 8 hours later. Altogether, these transcriptional regulations pointed to transient and complex modulations of ABA signaling taking place during the early germination of *pme36-1*.

Interestingly, auxin signaling also appeared to be modulated in the mutant. First, *JUMONJI13* (*JMJ13*), encoding a histone H3K27me3 demethylase, which was recently shown to regulate flowering time (Zheng *et al*., 2019) and self-fertility in Arabidopsis (Keyzor *et al*., 2021), was upregulated and then downregulated in *pme36-1* seeds at 16 and 24 HPS, respectively. H3K27me3 is a repressive posttranslational histone modification which exerts epigenetic control of genes from the entire IAA signaling pathway (Lafos *et al*., 2011; He *et al*., 2012). By relieving the H3K27me3 repressive mark, *JMJ13* is therefore likely to impact IAA signaling and its transcriptional modulation in *pm36-1* might affect the hormone signaling during germination of the mutant. Second, *CYP79B2* was upregulated in 16 HPS *pme36-1* seeds. This gene encodes a key enzyme involved in the conversion of L-tryptophan (Trp) to IAOx (indole-3-acetaldoxime), which is a common intermediate in Trp-dependent auxin biosynthesis pathway (reviewed in Casanova-Sáez et al. (2021)). Increase in IAOx levels result in elevated IAA biosynthesis and, thus, upregulation of *CYP79B2* in 16 HPS *pme36-1* seeds suggested that overproduction of the hormone arose in the mutant during seed imbibition. Nevertheless, IAOx is also a well-known precursor of indole glucosynolates (IGs), in a biosynthetic pathway for which the first step involves the *SUR2* gene, encoding the cytochrome P450 CYP83B1 protein (Barlier *et al*., 2000; Bak & Feyereisen, 2001). This gene was precisely also upregulated in 16 HPS *pme36-1* seed, thereby probably easing the potential IAA overproduction in the mutant seed to the benefit of the biosynthesis of IGs. Lastly, two genes encoding AUX/IAA transcriptional repressors, *IAA2* and *IAA33*, were downregulated in 16 HPS *pme36-1* seed. AUX/IAA proteins are auxin co-receptors and their degradation is crucial for auxin action (reviewed in Salehin et al. (2015)). Recently, IAA33 was shown to be a non-canonical AUX/IAA protein negatively regulating auxin signaling through competition with canonical AUX/IAA repressor IAA5 (Lv *et al*., 2020).

Overall, our results showed that *pme36-1* seed displayed a highly dynamic transcriptional regulation at the outset of germination. As several transcriptional regulations related to ABA and auxin signaling were highlighted in the mutant, further investigations were conducted for the analysis of homeostasis and the role of these hormones during *pme36-1* seed-to-seedling transition.

### *pme36-1* mature seeds contain reduced levels of ABA and IAA but display the same hormone sensitivity

ABA plays a major function in the regulation of dormancy and germination (reviewed in Shu et al. (2016)). Interestingly, during seed germination ABA treatment was shown to counteract the expected decline of PME activity by maintaining its high level beyond the testa rupture stage (Müller *et al*., 2013). In addition, the effect of the hormone was less efficient in delaying endosperm rupture in *PMEI5* overexpressing seeds, where DM of pectin remained high due to PME activity inhibition. It was thus hypothesized that PME was involved in the ABA-dependent temporal regulation of endosperm rupture. However, the interplay between ABA and the DM of pectin in the control of seed germination remains poorly documented. To gain insight into this connection, we investigated the role of ABA during *pme36-1* seed germination. First, ABA quantification was performed in mature seeds as well as in 24 HPS and 48 HPS seedlings from Col-0 and *pme36-1* lines. Interestingly, a strong reduction in ABA content was found in *pme36-1* mature seeds compared to the wild type (**Fig. 4A**). After germination, this difference decreased in the mutant, recovering to the wild-type ABA level in 48 HPS *pme36-1* seedlings. The reduction in ABA content, at the outset of germination in the mutant, explained the decrease in ABA signaling suggested by the down-regulation of *PYL13* expression as well as specific transcriptional regulations observed in 16 HPS mutant seeds, as previously detailed (**Dataset S1**). To assess whether the sensitivity to ABA remained effective in the mutant, we analyzed the effect of hormone supplementation on *pme36-1* and Col-0 germination in non-stratified seeds (**Fig. 4B**). As expected, ABA treatment substantially delayed endosperm rupture in the wild type, but the same delay was observed in the mutant, indicating the hormone sensitivity was not affected in *pme36-1* seeds. The decrease of the ABA level in *pme36-1* mature seeds would be expected to constitute an effective signal, which should trigger early germination in mutant seeds. This effect should have been even more efficient due to its additive combination with the one expected from the high DM of pectin, inducing release of dormancy and premature germination in *pme36-1* seed. But this was absolutely not the case, with *pme36-1* following a wild-type germination kinetic, which added a new level of complexity in the crosstalk between ABA homeostasis regulation and pectin methylesterification.

**Figure 4.**
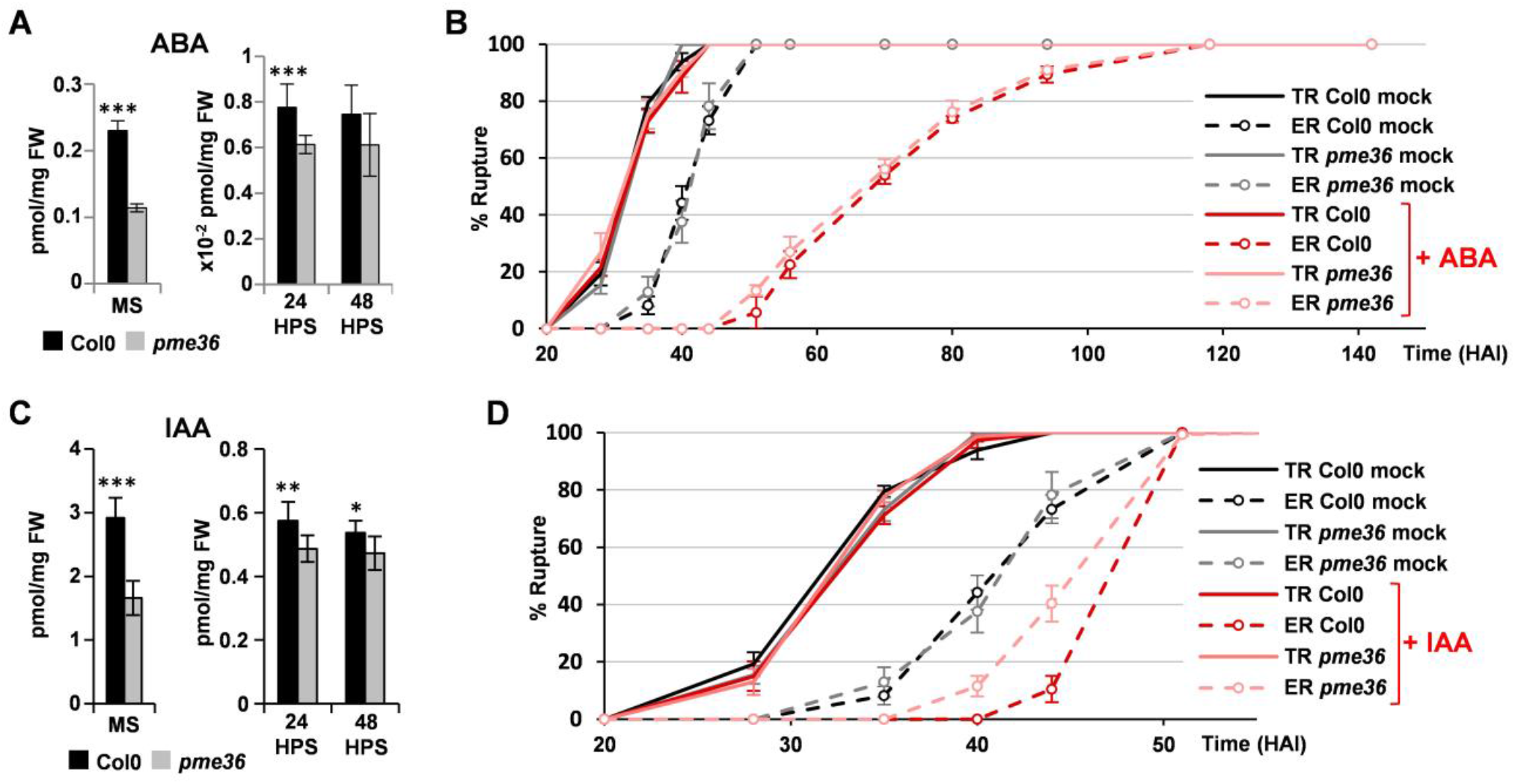
ABA and IAA homeostasis was strongly impaired although hormone sensitivity was not altered in *pme36-1* seed. **(A)** Quantification of ABA in *pme36-1* and Col-0 mature seed (MS) and in 24 and 48 HPS seedlings. Error bars indicate +/-SD of nine biological replicates. ANOVA comparison posttest indicated that the values indicated by asterisks were significantly different between wild-type and mutant: (*) for P<0.05; (**) for P<0.01; (***) for P<0.001; n=9. FW: fresh weight. **(B)** Effect of ABA treatment on germination kinetic of *pme36-1* and Col-0 seeds, characterized by testa rupture (TR) and endosperm rupture (ER). Data represent averages and error bars indicate +/- SE, obtained from three independent biological replicates of at least 50 seeds each. **(C)** Quantification of free IAA in the same conditions as in (A). **(D)** Effect of IAA treatment on germination kinetic of *pme36-1* and Col-0 seeds, analyzed as in (B).

While ABA and GA are classically considered as the main hormones that antagonistically regulate seed dormancy and germination, over the past few years auxin has emerged as new master player acting in crosstalk with ABA (reviewed in Shu et al. (2016) and Matilla (2020)). To evaluate whether auxin homeostasis was impacted by the modification of pectin methylesterification in *pme36-1*, IAA quantification was performed in seeds and seedlings from *pme36-1* and Col-0. Surprisingly, a strong decrease in IAA content was quantified in *pme36-1* mature seeds (**Fig. 4C**). After germination, the difference in IAA content between the mutant and the wild type decreased and remained negligible in 48 HPS seedlings. To determine whether the IAA decrease could affect the germination process in the mutant, we assessed *pme36-1* and Col-0 germination under IAA supplementation (**Fig. 4D**). IAA treatment did not affect the testa rupture and only slightly delayed the endosperm rupture in Col-0 and in *pme36-1*. The effect of IAA supplementation was less pronounced in the mutant, which makes sense considering the lower IAA content in the mutant, making probably the small effect produced by the supplementation less efficient in the mutant than in the wild type. The subtle negative effect of auxin on germination indicated that the IAA decrease observed in *pme36-1* mature seeds might have either no effect on germination or only a small positive one. Either way, this effect was not comparable with the strong activation the decrease in ABA content was expected to exert on the germination of the mutant.

### Auxin homeostasis is modulated through increase of IAA conjugation in *pme36-1* seed

The strong reduction of the IAA level in the mutant seeds and its subsequent fast recovery once germination had begun, were intriguing and suggested that a very specific regulation of auxin homeostasis occurred in the mutant. Conjugation of various amino acids to IAA, leading to inactivation or degradation, is a common way by which auxin homeostasis is modified, especially when dynamic and reversible modification of IAA concentration is required (Mellor *et al*., 2016; Casanova-Sáez *et al*., 2021). We quantified the level of two major IAA inactive conjugates, IAGlu (N-(Indole-3-ylAcetyl)-Glutamate) and IAAsp (N-(Indole-3-ylAcetyl)-Aspartate), and the level of the oxidative form oxIAA that induces IAA degradation, in mature seeds as well as in 24 and 48 HPS seedlings from Col-0 and *pme36-1* lines (**Fig. 5A,B,C**). Interestingly, the levels of IAGlu, IAAsp and oxIAA were strongly increased in *pme36-1* mature seeds and rapidly returned to the wild-type level during germination, except for IAGlu which remained at a higher level in the mutant seedlings compared to the wild type. These results indicated that the decrease of free IAA level observed in *pme36-1* mature seeds and the fast recovery to the wild-type level during germination involved specific regulation of conjugate production and IAA degradation, highlighting the dynamic regulation of the auxin homeostasis occurring during the seed-to-seedling transition in the mutant. Auxin conjugation is catalyzed by acyl-acid-amido synthetases encoded by auxin-induced *GRETCHEN HAGEN 3* (*GH3*) genes belonging to a family of 20 genes in Arabidopsis (Staswick *et al*., 2005). We quantified the expression of the *GH3* genes by RT-qPCR in *pme36-1* mature seed (**Fig. 5D**). While *GH3*.*7* transcript content decreased, the quantity of *GH3*.*3, GH3*.*4, GH3*.*5* and *GH3*.*6* transcripts strongly increased in mature seeds of the mutant compared to the wild type, suggesting that increased IAA conjugation was sustained by transcriptional upregulation of these *GH3* genes during *pme36-1* seed maturation. The expression of *GH3*.*3, GH3*.*5* and *GH3*.*6* genes were previously shown to be regulated by the *AUXIN RESPONSE FACTOR* genes *ARF6* and *ARF8*, as positive regulators, and *ARF17*, as negative regulator (Gutierrez *et al*., 2012). Interestingly, the expression of these 3 *ARFs* increased in 13 DPA *pme36-1* siliques, *ARF6* and *ARF8* displaying a stronger upregulation than *ARF17*, thereby providing a positive balance for subsequent activation of *GH3* gene expression (**Fig 5E**). It is noteworthy that, *PME36* expression increasing from 10 to 13 DPA siliques in the wild type, the defect in PME activity in the mutant cells was presumably perceived within this interval, which means that the gene expression modulation of the 3 *ARFs* in 13 DPA siliques was a very fast event. Therefore, the modulation of auxin homeostasis in *pme36-1* seed, through transcriptomic regulation of the ARF/GH3 module, constituted an early response to *PME36* disruption during seed maturation, occurring before the implementation of the compensatory *PME* and *PMEI* transcriptomic regulation. This most probably led to the decreased auxin content at the same time as the pectin methylesterification pattern was disrupted in the mutant seed. This concomitance suggested that the reduction of auxin content in *pme36-1* seeds might be involved in setting up the transcriptomic *PME*/*PMEI* compensatory mechanism.

**Figure 5.**
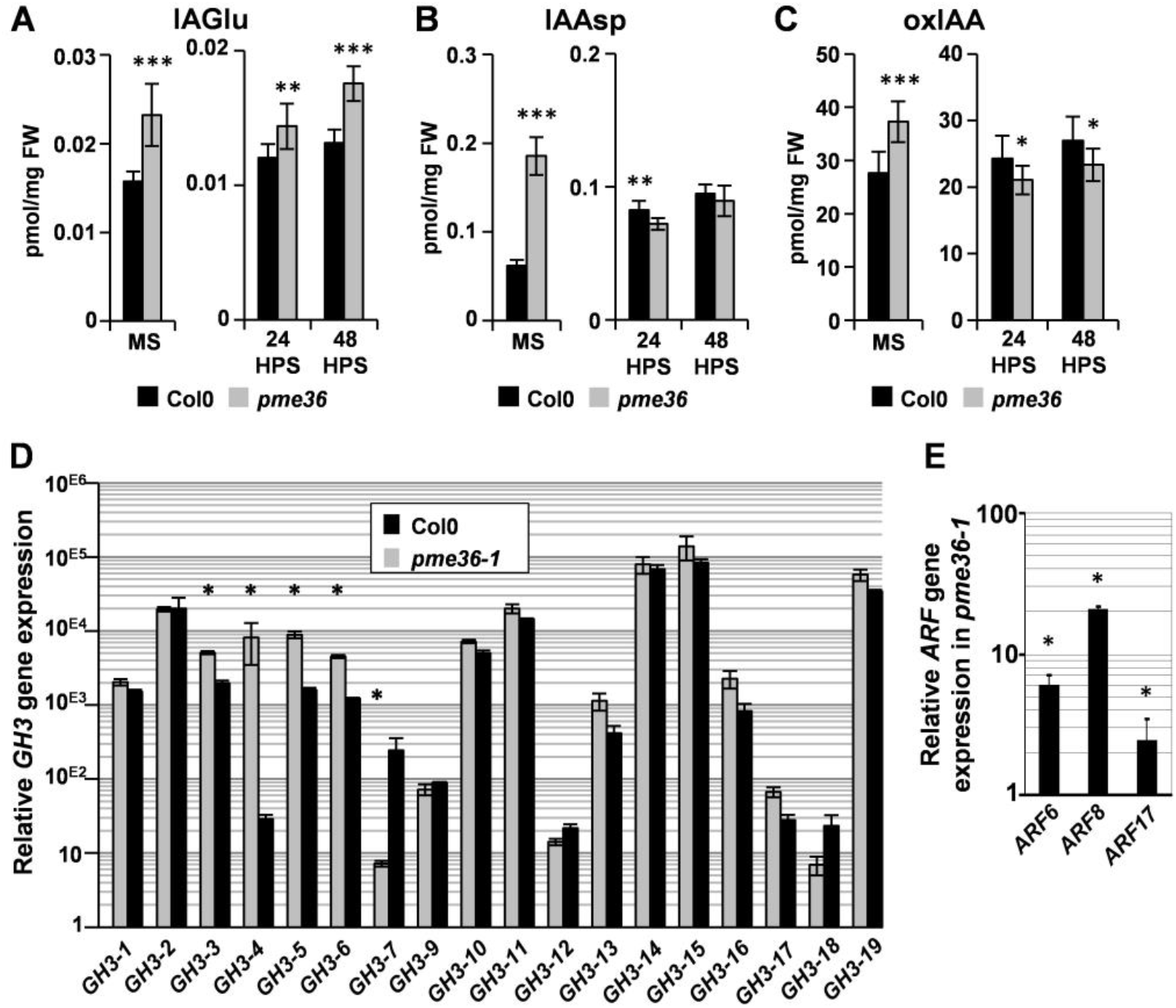
Auxin homeostasis is modulated through increase of IAA conjugation in *pme36-1* seed. **(A)** to **(C)** Quantification of IAGlu (N-(Indole-3-ylAcetyl)-Glutamate) and IAAsp (N-(Indole-3-ylAcetyl)-Aspartate) conjugates, and oxIAA catabolite, in the same conditions as in Fig. 4A. **(D)** Quantification of *GH3* transcripts by qRT-PCR in *pme36-1* and Col0 mature seed. Gene expression values shown are relative to the expression of TIP41, which has been validated as reference gene in our experimental conditions (see Methods). Error bars indicate +/- SE obtained from three independent qPCR experiments. The analysis was performed in two additional independent biological replicates, which gave similar results. A one-way analysis of variance combined with the Tukey’s multiple comparison posttest confirmed that the differences between the wild type and the mutants are significant; (*) P<0.05, n=3. No transcript was detected for GH3-8 and GH3-20. **(E)** Quantification of *ARF6, ARF8* and *ARF17* transcripts by qRT-PCR in *pme36-1* and Col-0 13 DPA siliques, analyzed as in (H).

### Transcriptomic and hormonal regulations are triggered by pectin remodeling in a dynamic homeostatic system ensuring the seed-to-seedling transition

Homeostatic systems are not easy to characterize since their main function is precisely to make silent transient variations that might occur during the life of an organism. When variations are too drastic, as for epistatic effects of a steady gene overexpression, homeostatic systems failed to maintain a physiological equilibrium, which most often leads to developmental defects. In this study, we took advantage of the strong, but transient, effect *PME36* disruption had on pectin methylesterification in the mature seed. By analyzing the physiological perturbations induced in the mutant, in the absence of growth phenotypical effects, we unveiled the complexity of an undiscovered homeostatic system likely ensuring the maintenance of the seed-to-seedling transition. Connections between the different regulatory pathways, deduced from modulations observed in *pme36-1*, are recapitulated in **Fig. 6** where consecutive steps are numbered. Our work clearly shows that, beyond its mechanical role within the cell wall, pectin methylesterification status acts as an upstream modulator of diverse regulatory pathways involved in plant growth and development. With this in mind, effects of cell wall-related mutations should be analyzed as consequences of both the straightforward change in cell wall properties and also the modulation of signaling pathways impacting diverse regulatory networks.

**Figure 6.**
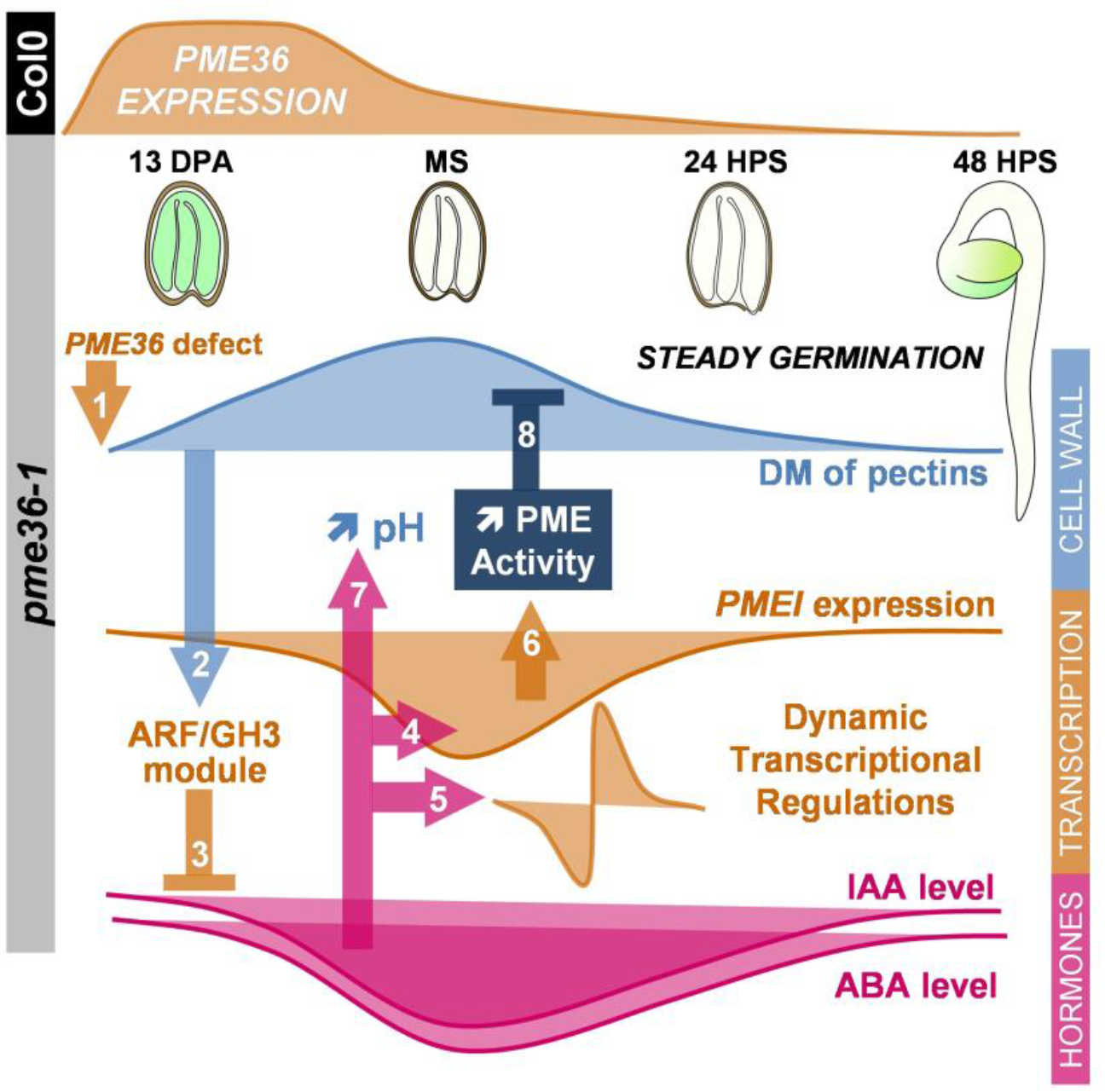
Transcriptomic and hormonal regulations are triggered by pectin remodeling in a dynamic homeostatic system ensuring the seed-to-seedling transition. First, *PME36* impairment negatively impacts on PME activity during seed maturation, which leads to an increase in the DM of pectins [1]. This triggers transcriptional activation of the ARF/GH3 module [2], leading to the reduction of IAA levels through increased conjugation activity [3]. Simultaneously, ABA content also decreases, which is possibly linked to the IAA level reduction given the connections previously shown between the signaling pathways of these two hormones during seed maturation and germination (Belin *et al*., 2009; Liu *et al*., 2013; Pellizzaro *et al*., 2020). Given the role of auxin in the regulation of cell wall-related genes (Majda & Robert, 2018), hormonal deregulation is likely to trigger subsequent transcriptional regulations in *pme36-1* (i) on *PME* and *PMEI* genes during seed maturation [4] and (ii) on a wider set of genes during early germination [5], among which some genes related to hormone signaling. The drop in *PMEI* transcript level in mutant seeds is likely to reduce the quantity of neo-synthesized PMEI proteins during the imbibition stage, thus releasing PME inhibition at the outset of germination [6]. Concomitantly, the reduction in IAA levels, known to mediate cell wall acidification which triggers acid growth (Fendrych *et al*., 2016), probably leads to a transient increase of apoplast pH in imbibed mutant seeds [7]. This might constitute an additive positive effect, reducing PMEI-PME interactions for which stability declines when pH value rises to neutrality and, at the same time, initiating PME activity, which has an optimum at alkaline pH (Sénéchal *et al*., 2015; Hocq *et al*., 2017a; Hocq *et al*., 2017b; Sénéchal *et al*., 2017). Moreover, a pH increase might also result in the reduction of pH-dependent activity of wall-loosening agents like expansins (Cosgrove, 2016), thereby counterbalancing the potential loosening effect - expected to promote germination acceleration - the high DM of pectins might cause in *pme36-1* imbibed seeds and avoiding wall-loosening reactivation during the pectin de-methylesterification process in germinating mutant seeds. Finally, enhanced PME activity allows a rapid decrease of the DM of pectin to wild-type levels in *pme36-1* young seedlings [8]. Overall, all these factors (modulation of the DM of pectins, transcriptome dynamics and hormone homeostasis) appeared to be accurately regulated to generate a physiological equilibrium that allows normal germination of the mutant seed. This sheds light on a previously undescribed regulatory network integrating different pathways, connected to pectin methylesterification status, to constitute a homeostatic system ensuring the seed-to-seedling transition.

## ACKNOWLEDGMENTS

This work was supported by two grants from the Agence Nationale de la Recherche (ANR-09-BLANC-0007-01 GROWPEC project and ANR-14-CE34-0010 PECTOSIGN project), by the Conseil Régional des Hauts-de-France and by the Université de Picardie Jules Verne. J.P. acknowledges the funding from the Institut Universitaire de France (IUF). K.J.D.L. and J.P.K. acknowledge funding from the UK Biotechnology and Biosciences Research Council (BB/G024898/1) through the ERA-NET support of the vSeed programme. This work was also funded by the Ministry of Education, Youth and Sports of the Czech Republic (European Regional Development Fund-Project “Plants as a tool for sustainable global development” No. CZ.02.1.01/0.0/0.0/16_019/0000827).

## AUTHOR CONTRIBUTIONS

F.J. and S.G. contributed equally to this work. F.J., S.B. and L.G. developed the RT-qPCR assay on *PME* and *PMEI* genes, with the help of S.G., F.S., L.H. and Ga.M.; S.G., Ga.M, H.D. and L.G. performed the RT-qPCR analysis, the GUS assays, the PME activity quantification and the germination testing. S.G. and L.G. performed the microarrays analysis. P.A., M.S. and O.N. performed the hormone measurements. A.V., S.P. and S.V. developed and performed the enzymatic fingerprinting of pectins. K.J.D.L. and J.P.K. performed the immunofluorescence microscopy. Gr.M., S.G., F.J., J.P. and L.G. designed the research and analyzed the data. L.G. and J.P. conceptualized and supervised the overall project. L.G. wrote the article with inputs from all co-authors. J.P.K. edited the article.

## SUPPORTING INFORMATION

Figure S1. *pme36-1* and *pme36-2* are knockout mutants for *PME36* gene.

Table S1. Sequences of primers designed for RT-qPCR experiment on *PME* and *PMEI* genes.

Dataset S1. Genes differentially expressed in 16 and 24 HPS *pme36-1* seedlings compared to Col-0.

## REFERENCES

Andres-Robin A, Reymond MC, Brunoud G, Martin-Magniette ML, Monéger F, Scutt CP. 2020. Immediate targets of ETTIN suggest a key role for pectin methylesterase inhibitors in the control of Arabidopsis gynecium development. Plant Signaling and Behavior 15.

Andres-Robin A, Reymond MC, Dupire A, Battu V, Dubrulle N, Mouille G, Lefebvre V, Pelloux J, Boudaoud A, Traas J, et al. 2018. Evidence for the regulation of gynoecium morphogenesis by ETTIN via cell wall dynamics. Plant Physiology 178: 1222–1232.

Arvidsson S, Kwasniewski M, Riaño-Pachón DM, Mueller-Roeber B. 2008. QuantPrime – a flexible tool for reliable high-throughput primer design for quantitative PCR. BMC Bioinformatics 2008 9:1 9: 1–15.

Bai B, Van Der Horst S, Cordewener JHG, America TAHP, Hanson J, Bentsink L. 2020. Seed-stored mRNAs that are specifically associated to monosomes are translationally regulated during germination. Plant Physiology 182: 378–392.

Bak S, Feyereisen R. 2001. The involvement of two P450 enzymes, CYP83B1 and CYP83A1, in auxin homeostasis and glucosinolate biosynthesis. Plant Physiology 127: 108–118.

Barlier I, Kowalczyk M, Marchant A, Ljung K, Bhalerao R, Bennett M, Sandberg G, Bellini C. 2000. The SUR2 gene of Arabidopsis thaliana encodes the cytochrome P450 CYP83B1, a modulator of auxin homeostasis. Proceedings of the National Academy of Sciences of the United States of America 97: 14819–14824.

Belin C, Megies C, Hauserová E, Lopez-Molina L. 2009. Abscisic acid represses growth of the arabidopsis embryonic axis after germination by enhancing auxin signalings. Plant Cell 21: 2253–2268.

Braybrook SA, Peaucelle A. 2013. Mechano-Chemical Aspects of Organ Formation in Arabidopsis thaliana: The Relationship between Auxin and Pectin. PLoS ONE 8.

Casanova-Sáez R, Mateo-Bonmatí E, Ljung K. 2021. Auxin metabolism in plants. Cold Spring Harbor Perspectives in Medicine 11: 1–23.

Chebli Y, Bidhendi AJ, Kapoor K, Geitmann A. 2021. Cytoskeletal regulation of primary plant cell wall assembly. Current Biology 31: R681–R695.

Chen H, Ruan J, Chu P, Fu W, Liang Z, Li Y, Tong J, Xiao L, Liu J, Li C, et al. 2020. AtPER1 enhances primary seed dormancy and reduces seed germination by suppressing the ABA catabolism and GA biosynthesis in Arabidopsis seeds. Plant Journal 101: 310–323.

Clausen MH, Willats WGT, Knox JP. 2003. Synthetic methyl hexagalacturonate hapten inhibitors of anti-homogalacturonan monoclonal antibodies LM7, JIM5 and JIM7. Carbohydrate Research 338: 1797–1800.

Cosgrove DJ. 2016. Catalysts of plant cell wall loosening. F1000Research 5.

Cosgrove DJ. 2018. Diffuse growth of plant cell walls. Plant Physiology 176: 16–27.

Dekkers BJW, Pearce S, van Bolderen-Veldkamp RP, Marshall A, Widera P, Gilbert J, Drost HG, Bassel GW, Müller K, King JR, et al. 2013. Transcriptional dynamics of two seed compartments with opposing roles in Arabidopsis seed germination. Plant Physiology 163: 205–215.

Du J, Kirui A, Huang S, Wang L, Barnes WJ, Kiemle SN, Zheng Y, Rui Y, Ruan M, Qi S, et al. 2020. Mutations in the Pectin Methyltransferase QUASIMODO2 influence cellulose biosynthesis and wall integrity in arabidopsis. Plant Cell 32: 3576–3597.

Fendrych M, Leung J, Friml J. 2016. Tir1/AFB-Aux/IAA auxin perception mediates rapid cell wall acidification and growth of Arabidopsis hypocotyls. eLife 5.

Floková K, Tarkowská D, Miersch O, Strnad M, Wasternack C, Novák O. 2014. UHPLC-MS/MS based target profiling of stress-induced phytohormones. Phytochemistry 105: 147–157.

Fuchs S, Tischer S V., Wunschel C, Christmann A, Grill E. 2014. Abscisic acid sensor RCAR7/PYL13, specific regulator of protein phosphatase coreceptors. Proceedings of the National Academy of Sciences of the United States of America 111: 5741–5746.

Guénin S, Mareck A, Rayon C, Lamour R, Assoumou Ndong Y, Domon JM, Sénéchal F, Fournet F, Jamet E, Canut H, et al. 2011. Identification of pectin methylesterase 3 as a basic pectin methylesterase isoform involved in adventitious rooting in Arabidopsis thaliana. New Phytologist 192: 114–126.

Gutierrez L, Bussell JD, Pǎcurar DI, Schwambach J, Pǎcurar M, Bellini C. 2009. Phenotypic plasticity of adventitious rooting in arabidopsis is controlled by complex regulation of AUXIN RESPONSE FACTOR transcripts and microRNA abundance. Plant Cell 21: 3119–3132.

Gutierrez L, Conejero G, Castelain M, Guénin S, Verdeil JL, Thomasset B, Van Wuytswinkel O. 2006. Identification of new gene expression regulators specifically expressed during plant seed maturation. In: Journal of Experimental Botany. J Exp Bot, 1919–1932.

Gutierrez L, Mongelard G, Floková K, Pǎcurar DI, Novák O, Staswick P, Kowalczyk M, Pǎcurar M, Demailly H, Geiss G, et al. 2012. Auxin controls Arabidopsis adventitious root initiation by regulating jasmonic acid homeostasis. Plant Cell 24: 2515–2527.

He C, Chen X, Huang H, Xu L. 2012. Reprogramming of H3K27me3 Is Critical for Acquisition of Pluripotency from Cultured Arabidopsis Tissues. PLoS Genetics 8.

Hocq L, Guinand S, Habrylo O, Voxeur A, Tabi W, Safran J, Fournet F, Domon JM, Mollet JC, Pilard S, et al. 2020. The exogenous application of AtPGLR, an endo-polygalacturonase, triggers pollen tube burst and repair. Plant Journal 103: 617–633.

Hocq L, Pelloux J, Lefebvre V. 2017a. Connecting Homogalacturonan-Type Pectin Remodeling to Acid Growth. Trends in Plant Science 22: 20–29.

Hocq L, Sénéchal F, Lefebvre V, Lehner A, Domon JM, Mollet JC, Dehors J, Pageau K, Marcelo P, Guérineau F, et al. 2017b. Combined experimental and computational approaches reveal distinct pH dependence of pectin Methylesterase Inhibitors. Plant Physiology 173: 1075–1093.

Jonsson K, Lathe RS, Kierzkowski D, Routier-Kierzkowska AL, Hamant O, Bhalerao RP. 2021. Mechanochemical feedback mediates tissue bending required for seedling emergence. Current Biology 31: 1154–1164.e3.

Keyzor C, Mermaz B, Trigazis E, Jo SY, Song J. 2021. Histone Demethylases ELF6 and JMJ13 Antagonistically Regulate Self-Fertility in Arabidopsis. Frontiers in Plant Science 12.

Lafos M, Kroll P, Hohenstatt ML, Thorpe FL, Clarenz O, Schubert D. 2011. Dynamic regulation of H3K27 trimethylation during arabidopsis differentiation. PLoS Genetics 7.

Lee KJD, Dekkers BJW, Steinbrecher T, Walsh CT, Bacic A, Bentsink L, Leubner-Metzger G, Paul Knox J. 2012. Distinct cell wall architectures in seed endosperms in representatives of the Brassicaceae and Solanaceae. Plant Physiology 160: 1551–1566.

Lefebvre V, North H, Frey A, Sotta B, Seo M, Okamoto M, Nambara E, Marion-Poll A. 2006. Functional analysis of Arabidopsis NCED6 and NCED9 genes indicates that ABA synthesized in the endosperm is involved in the induction of seed dormancy. Plant Journal 45: 309–319.

Levesque-Tremblay G, Müller K, Mansfield SD, Haughn GW. 2015a. Highly methyl esterified seeds is a pectin methyl esterase involved in embryo development. Plant Physiology 167: 725–737.

Levesque-Tremblay G, Pelloux J, Braybrook SA, Müller K. 2015b. Tuning of pectin methylesterification: consequences for cell wall biomechanics and development. Planta 242: 791–811.

Liu X, Zhang H, Zhao Y, Feng Z, Li Q, Yang HQ, Luan S, Li J, He ZH. 2013. Auxin controls seed dormancy through stimulation of abscisic acid signaling by inducing ARF-mediated ABI3 activation in Arabidopsis. Proceedings of the National Academy of Sciences of the United States of America 110: 15485–15490.

Lv B, Yu Q, Liu J, Wen X, Yan Z, Hu K, Li H, Kong X, Li C, Tian H, et al. 2020. Non-canonical AUX / IAA protein IAA 33 competes with canonical AUX / IAA repressor IAA 5 to negatively regulate auxin signaling. The EMBO Journal 39.

Majda M, Robert S. 2018. The role of auxin in cell wall expansion. International Journal of Molecular Sciences 19.

Martinez DE, Borniego ML, Battchikova N, Aro EM, Tyystjärvi E, Guiamét JJ. 2015. SASP, a Senescence-Associated Subtilisin Protease, is involved in reproductive development and determination of silique number in Arabidopsis. Journal of Experimental Botany 66: 161–174.

Matilla AJ. 2020. Auxin: Hormonal signal required for seed development and dormancy. Plants 9: 1–17.

Mellor N, Band LR, Pěňík A, Novák O, Rashed A, Holman T, Wilson MH, Vo U, Bishopp A, King JR, et al. 2016. Dynamic regulation of auxin oxidase and conjugating enzymes AtDAO1 and GH3 modulates auxin homeostasis. Proceedings of the National Academy of Sciences of the United States of America 113: 11022–11027.

Müller K, Levesque-Tremblay G, Bartels S, Weitbrecht K, Wormit A, Usadel B, Haughn G, Kermode AR. 2013. Demethylesterification of cell wall pectins in Arabidopsis plays a role in seed germination. Plant Physiology 161: 305–316.

Ogawa M, Kay P, Wilson S, Swain SM. 2009. Arabidopsis Dehiscence Zone Polygalacturonase1 (ADPG1), ADPG2, and Quartet2 are polygalacturonases required for cell separation during reproductive development in Arabidopsis. Plant Cell 21: 216–233.

Pellizzaro A, Neveu M, Lalanne D, Ly Vu B, Kanno Y, Seo M, Leprince O, Buitink J. 2020. A role for auxin signaling in the acquisition of longevity during seed maturation. New Phytologist 225: 284–296.

Rajjou L, Gallardo K, Debeaujon I, Vandekerckhove J, Job C, Job D. 2004. The effect of α-amanitin on the arabidopsis seed proteome highlights the distinct roles of stored and neosynthesized mRNAs during germination. Plant Physiology 134: 1598–1613.

Salehin M, Bagchi R, Estelle M. 2015. ScfTIR1/AFB-based auxin perception: Mechanism and role in plant growth and development. Plant Cell 27: 9–19.

Sano N, Rajjou L, North HM. 2020. Lost in Translation: Physiological Roles of Stored mRNAs in Seed Germination. Plants (Basel, Switzerland) 9.

Scheler C, Weitbrecht K, Pearce SP, Hampstead A, Büttner-Mainik A, Lee KJD, Voegele A, Oracz K, Dekkers BJW, Wang X, et al. 2015. Promotion of testa rupture during garden cress germination involves seed compartment-specific expression and activity of pectin methylesterases. Plant Physiology 167: 200–215.

Schoenaers S, Balcerowicz D, Breen G, Hill K, Zdanio M, Mouille G, Holman TJ, Oh J, Wilson MH, Nikonorova N, et al. 2018. The Auxin-Regulated CrRLK1L Kinase ERULUS Controls Cell Wall Composition during Root Hair Tip Growth. Current Biology 28: 722–732.e6.

Sechet J, Frey A, Effroy-Cuzzi D, Berger A, Perreau F, Cueff G, Charif D, Rajjou L, Mouille G, North HM, et al. 2016. Xyloglucan metabolism differentially impacts the cell wall characteristics of the endosperm and embryo during Arabidopsis seed germination. Plant Physiology 170.

Sénéchal F, Habrylo O, Hocq L, Domon JM, Marcelo P, Lefebvre V, Pelloux J, Mercadante D. 2017. Structural and dynamical characterization of the pH-dependence of the pectin methylesterase-pectin methylesterase inhibitor complex. Journal of Biological Chemistry 292: 21538–21547.

Sénéchal F, L’Enfant M, Domon JM, Rosiau E, Crépeau MJ, Surcouf O, Esquivel-Rodriguez J, Marcelo P, Mareck A, Guérineau F, et al. 2015. Tuning of pectin methylesterification: Pectin methylesterase inhibitor 7 modulates the processive activity of co-expressed pectin methylesterase 3 in a pH-dependentmanner. Journal of Biological Chemistry 290: 23320–23335.

Shigeyama T, Watanabe A, Tokuchi K, Toh S, Sakurai N, Shibuya N, Kawakami N. 2016. α-Xylosidase plays essential roles in xyloglucan remodelling, maintenance of cell wall integrity, and seed germination in Arabidopsis thaliana. Journal of Experimental Botany 67: 5615–5629.

Shu K, Liu XD, Xie Q, He ZH. 2016. Two Faces of One Seed: Hormonal Regulation of Dormancy and Germination. Molecular Plant 9: 34–45.

Staswick PE, Serban B, Rowe M, Tiryaki I, Maldonado MT, Maldonado MC, Suza W. 2005. Characterization of an arabidopsis enzyme family that conjugates amino acids to indole-3-acetic acid. Plant Cell 17: 616–627.

Tischer S V., Wunschel C, Papacek M, Kleigrewe K, Hofmann T, Christmann A, Grill E. 2017. Combinatorial interaction network of abscisic acid receptors and coreceptors from Arabidopsis thaliana. Proceedings of the National Academy of Sciences of the United States of America 114: 10280–10285.

Vaahtera L, Schulz J, Hamann T. 2019. Cell wall integrity maintenance during plant development and interaction with the environment. Nature Plants 5: 924–932.

Verhertbruggen Y, Marcus SE, Haeger A, Ordaz-Ortiz JJ, Knox JP. 2009. An extended set of monoclonal antibodies to pectic homogalacturonan. Carbohydrate Research 344: 1858–1862.

Vlad F, Rubio S, Rodrigues A, Sirichandra C, Belin C, Robert N, Leung J, Rodriguez PL, Laurière C, Merlot S. 2009. Protein phosphatases 2C regulate the activation of the Snf1-related kinase OST1 by abscisic acid in Arabidopsis. Plant Cell 21: 3170–3184.

Voiniciuc C, Dean GH, Griffiths JS, Kirchsteiger K, Hwang YT, Gillett A, Dow G, Western TL, Estelle M, Haughn GW. 2013. Flying saucer1 is a transmembrane RING E3 ubiquitin ligase that regulates the degree of pectin methylesterification in Arabidopsis seed mucilage. Plant Cell 25: 944–959.

Voxeur A, Habrylo O, Guénin S, Miart F, Soulié MC, Rihouey C, Pau-Roblot C, Domon JM, Gutierrez L, Pelloux J, et al. 2019. Oligogalacturonide production upon Arabidopsis thaliana-Botrytis cinerea interaction. Proceedings of the National Academy of Sciences of the United States of America 116: 19743–19752.

Wang Q, Guo Q, Guo Y, Yang J, Wang M, Duan X, Niu J, Liu S, Zhang J, Lu Y, et al. 2018. Arabidopsis subtilase SASP is involved in the regulation of ABA signaling and drought tolerance by interacting with OPEN STOMATA 1. Journal of Experimental Botany 69: 4403–4417.

Wang H, Wang H. 2015. Multifaceted roles of FHY3 and FAR1 in light signaling and beyond. Trends in Plant Science 20: 453–461.

Wang M, Yuan D, Gao W, Li Y, Tan J, Zhang X. 2013. A Comparative Genome Analysis of PME and PMEI Families Reveals the Evolution of Pectin Metabolism in Plant Cell Walls. PLoS ONE 8.

Willats WGT, Limberg G, Buchholt HC, Van Alebeek GJ, Benen J, Christensen TMIE, Visser J, Voragen A, Mikkelsen JD, Knox JP. 2000. Analysis of pectic epitopes recognised by hybridoma and phage display monoclonal antibodies using defined oligosaccharides, polysaccharides, and enzymatic degradation. Carbohydrate Research 327: 309–320.

Willats WGT, Orfila C, Limberg G, Buchholt HC, Van Alebeek GJWM, Voragen AGJ, Marcus SE, Christensen TMIE, Mikkelsen JD, Murray BS, et al. 2001. Modulation of the degree and pattern of methyl-esterification of pectic homogalacturonan in plant cell walls: Implications for pectin methyl esterase action, matrix properties, and cell adhesion. Journal of Biological Chemistry 276: 19404–19413.

Wolf S. 2017. Plant cell wall signalling and receptor-like kinases. Biochemical Journal 474: 471–492.

Xu P, Chen H, Cai W. 2020. Transcription factor CDF4 promotes leaf senescence and floral organ abscission by regulating abscisic acid and reactive oxygen species pathways in Arabidopsis. EMBO reports 21.

Zheng S, Hu H, Ren H, Yang Z, Qiu Q, Qi W, Liu X, Chen X, Cui X, Li S, et al. 2019. The Arabidopsis H3K27me3 demethylase JUMONJI 13 is a temperature and photoperiod dependent flowering repressor. Nature Communications 10.

Zhu Q, Zhang J, Gao X, Tong J, Xiao L, Li W, Zhang H. 2010. The Arabidopsis AP2/ERF transcription factor RAP2.6 participates in ABA, salt and osmotic stress responses. Gene 457: 1–12.

