## Supplemental Figure S1 for "Pectin remodeling belongs to a homeostatic system and triggers transcriptomic and hormonal modulations"

A

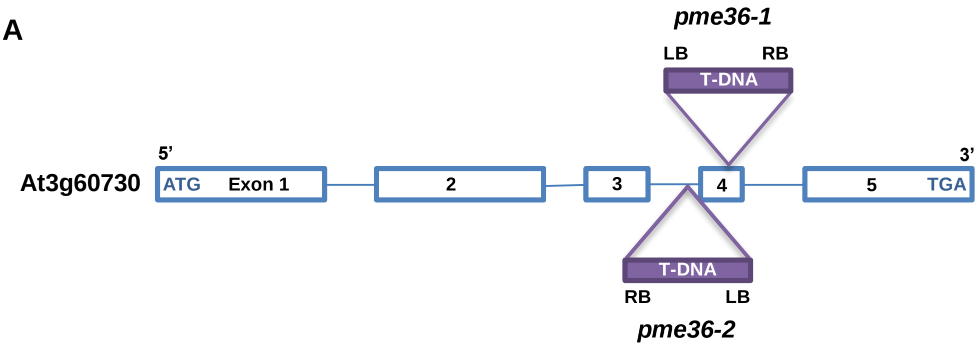


B

**At3g60730**

ATGTCCACGTTTGTAAAAGTCACCGACTTAATCACTATCATGTTCTTTCTAGCCATTGCTGCCGTCATAACCGCCTCAAACACAGCCGAG
TTGGATGTACTTGAAATGGCTAGAACTGCTGTGGTTGAAGCCAAGACTAGTTTTGGGTCAATGGCTGTGACTGAGGCAACTAGTGAAG
TTGCTGGTAGTTATTATAAATTGGGGTTAAGTGAATGCGAGAAGCTCTACGATGAGAGTGAAGCTAGGCTATCGAAACTGGTCGTAGAT
CATGAGAATTTCACCGTTGAAGATGTTAGAACGTGGCTAAGTGGCGTGCTTGCGAATCATCACACTTGTTTAGACGGTTTAATTCAGCA
ACGTCAAGGTCATAAGCCTTTGGTCCATAGCAACGTCACGTTCGTGCTCCATGAAGCTTTGGCTTTCTACAAAAAGTCTAGAGGTCATA
TGAAAAAGAgtaagaaaacttacacatacattcatgtatacataattaaataaaatgatacaataaatcgctctgttttatcactaaataacgattgtagactccaagataagtgtaaatttttgtttga
gatatcatggtttatctattatggatctctaataacatttggtagGGCTGCATGGACCAGCGAGACAGGGCCATGGACCAACGAGACCAAAACATAGACCAACG
AGACCAAATCATGGACCAGGAAGATCACACCATGGACCATCAAGACCAAACCAAAACGGTGGAATGTTAGTGTCATGGAATCCAACAA
GTTCAAGGGCAGATTTCGTGGTGGCTCGGGACGGTTCGGCGACTCATCGGACGATAAACCAAGCATTAGCCGCCGTTTCAAGAATGG
GTAAAAGTCGGTTGAATAGAGTGATAATATACATAAAAGCTGGTGTTTATAACGAAAAAATCGAAATAGATAGACACATGAAAAACATCA
TGCTAGTTGGTGATGGTATGGACCGTACAATTGTCACCAATAACCGAAACGTCCCGGACGGTTCCACGACTTATGGATCAGCGACATT
CGgtataaaatatttaaaattctactattaatcttcatacatagatcatgcatgttgaataattactgcgcttactcgttatatagGAGTGTCCGGGGATGGATTTTGGGCACGGGAC
ATAACGTTCGAGAATACTGCAGGACCACATAAACATCAAGCTGTGGCGTTGAGAGTGAGCTCAGACTTGTCTTTATTCTATCGGTGTAG
TTTCAAAGGATATCAAGATACTCTTTTCACGCACTCACTTCGTCAATTCTACCGCGATTGTCATATTTATGgtattatttttaatctatgtaccatttgtttaaa
ctcgtatggaagctagtcatgatcatacatggattaagaatttttgggaaattttggaatagGTACGATTGATTTCATATTTGGAGATGCGGCAGCTGTTTTTCAGAACT
GTGATATATTTGTGAGGCGGCCTATGGATCATCAAGGCAACATGATCACAGCTCAAGGAAGAGACGACCCACATACAAATTCAGgtaatgtt
ttgtttggccatcttttgactatcaAtcaggggtcatcaactataccgaaattattagaaccaaccgattctaatcagattatatttttaaaaa aattgttcatatgaaaatatccaaaaactaatgcatgcga
aagggaggaaaatcgaattaaaatttaagtgtaagtttgacttgataataatcaaacttgaacaaattaccaaaaaattaaaataatcaaacttgaacagaaatcgaattggaaaccaattttta acc
aaccccaaaccggaattcatattgaaaactcctatcgttaatagatacccatggttagtgttaaccatgaatttttttgtttaccttactggattagGCATTTCGATCCAACACTCACGGATC
AGGGCCGCACCAGAATTCGAAGCAGTTAAAGGCCGGTTTAAGAGCTACTTGGGCCGGCCATGGAAGAAGTACTCACGGACAGTGTTT
CTCAAAACAGATATCGACGAGTTGATTGACCCGAGAGGGTGGAGAGAATGGAGTGGCAGTTATGCTTTGTCGACGTTGTATTACGGCG
AGTTTATGAACACCGGAGCTGGAGCAGGGACTGGTAGAAGGGTGAACTGGCCGGGATTTCACGTCCTACGCGGTGAGGAAGAGGCTT
CTCCGTTTACCGTCAGCCGATTCATACAAGGAGATTCTTGGATCCCTATCACCGGTGTGCCGTTTTCAGCTGGGGTTTGA

acaaattacc : intronic sequence
AGCCGAT : exonic sequence
ATCGATC : annealing position of RT-PCR primers

**GA** : SALK_022170 insertion - *pme36-1*
**aa** : SALK_091350 insertion - *pme36-2*

C

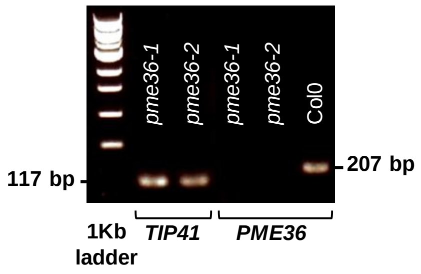


D

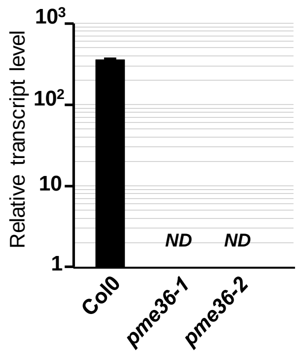


**Figure S1. *pme36-1* and *pme36-2* are knockout mutants for *PME36* gene.**
**(A)** Diagram showing the T-DNA insertion for each mutant. **(B)** Sequence of *PME36* transcript showing the site of each insertion and the annealing position of forward (ATCGGTGTAGTTTCAAAGG) and reverse (TTGTATGTGGGTCGTCTC) primers used in the RT-PCR experiment shown in (C). **(C)** RT-PCR performed on RNA extracted from seed, showing the absence of intact *PME36* RNA in both mutants. Production of amplicon from *TIP41* transcript was used to control the presence of cDNA in the samples from both mutants. Col0 was used to control the production of amplicon from *PME36* transcript. Electrophoresis was run in 1% agarose gel using 1Kb Ladder (Ozyme). (**D**) RT-qPCR confirming the absence of *PME36* transcript in the seeds from both mutants.
